# Validating the Standard Model of diffusion MRI in white matter with Numerical Substrates

**DOI:** 10.64898/2026.01.28.702302

**Authors:** Jasmine Nguyen-Duc, Quentin Uhl, Rita Oliveira, Jonathan Rafael-Patiño, Ileana O Jelescu

## Abstract

The non-invasive estimation of intra- and extracellular microstructural parameters using biophysical models has been a major focus in brain microstructure imaging with MRI. The Standard Model (SM) of diffusion in white matter (WM) provides a unifying framework for various modelling approaches, representing axons as impermeable narrow cylinders embedded within a locally anisotropic extra-axonal space. However, the SM relies on simplifying assumptions that may not hold in realistic WM tissue, as they do not take into account axonal undulations, beading, the presence of glial cells, or membrane permeability.

In this work, we investigate how SM-derived estimates behave when the model is applied to realistic numerical WM substrates generated by the CATERPillar tool. Specifically, we vary (i) axonal morphological features such as beading and undulations, (ii) axonal packing density, (iii) orientation dispersion, (iv) membrane permeability of axons and astrocytes separately, (v) myelin volume fraction, and (vi) diffusion time. In each part of the analysis, different noise levels are introduced. Overall, according to our results, the relative changes in SM estimates show that the intra-axonal volume fraction *f* increased with stronger beading, higher packing density, and greater myelin volume, and was strongly influenced by axonal and astrocytic permeability. The orientation dispersion index *p*_2_ was affected by undulation, but was substantially biased at low packing densities, with stronger beading and when astrocytes were impermeable. The effective intra-axonal diffusivity *D_a_* decreased with stronger beading and undulation and tended to be overestimated in most scenarios. The parallel extra-axonal diffusivity *D*_*e*∥_ was strongly influenced by axonal permeability, as well as packing density, dispersion, and undulation, and was the most noise-sensitive parameter, showing systematic overestimation at low SNR. Finally, the effective perpendicular extra-axonal diffusivity *D*_*e*⊥_ was the most stable parameter relative to the effective ground truth across the tested conditions, while remaining sensitive to packing density, axonal permeability, myelin volume fraction, and undulation. These findings enable users to identify potential biases introduced by varying conditions and to adjust their interpretations accordingly.

## 1 Introduction

Mapping tissue microstructure using diffusion magnetic resonance imaging (dMRI) has caught considerable attention from the scientific community. Its non-invasive application, view of intact *in situ* tissue, wide field of view and its relatively fast and cheap acquisition make it an excellent choice for studying the brain *in vivo* [1]. The underlying rationale is that dMRI signals reflect water diffusion within biological tissues, where diffusion trajectories are affected by cellular structures, such as cell membranes, making the signals sensitive to tissue microstructural organisation.

To investigate microstructural properties using dMRI, various approaches have been developed, with Diffusion Tensor Imaging (DTI) [2] being the most widely used. Its extension, diffusion kurtosis imaging (DKI) [3], captures non-Gaussian diffusion and provides additional metrics, yet both remain limited in specificity [4]. To overcome this, there has been an increasing interest in biophysical tissue models [5]. The forward problem consists of predicting diffusion signals analytically in simple known tissue geometries [6]. To solve the inverse problem, the analytical equations of the model, which describe relevant geometric and diffusion-related characteristics, are fit to diffusion signals [1, 7–9]. However, the inverse problem is inherently ill-posed as multiple tissue configurations can yield similar diffusion signals [10–12]. An additional complexity arises due to the relatively small size of the biological cells of interest (1-10 *µm*) compared to the typical voxel size (1-2 *mm* in width), causing the measured signal to represent an average over the entire voxel volume [6, 13]. This makes identifying specific microscopic length scales particularly difficult, as a huge amount of information is lost in this averaging [6, 13]. Consequently, biophysical models usually estimate compartment-specific diffusivities and volume fractions, which have “survived” the averaging, yet face challenges in accurately determining finer details.

The predominant multi-compartment biophysical model for white matter (WM) is known as the Standard Model (SM) [12, 14]. In this model, axons (and potentially glial processes) are depicted as impermeable, zero-radius cylinders, commonly termed “sticks”, arranged into locally coherent fibre bundles. Together, these axons form the intra-axonal space (IAS), with diffusivity that is assumed to only occur along each axon. On the other hand, diffusion in the extra-axonal space (EAS) of each locally coherent fibre bundle is represented by an axially symmetric diffusion tensor. Within each voxel, the fibres are distributed according to a fibre orientation distribution function (ODF), modelled up to its second order anisotropy invariant (*p*_2_). In total, the SM estimates five parameters: the axonal water fraction *f*, the ODF coefficient *p*_2_, the intra-axonal diffusion coefficient along the principal axonal orientation *D*_*a*_, the extra-axonal diffusion coefficient perpendicular to the principal axonal orientation *D*_*e*⊥_, and the extraaxonal diffusion coefficient parallel to the principal axonal orientation *D*_*e*∥_. It unifies multiple modelling frameworks [12, 14–29], all based on similar principles but differing in their assumptions [5, 7, 30].

The SM is widely accepted within the dMRI community as an appropriate model for WM [5, 31, 32], largely because myelinated axons are considered the dominant structural components. Despite its usefulness, the SM and biophysical models in general still fall short of capturing the true complexity of brain microstructure. While simplifications are necessary to limit the number of parameters to fit, some assumptions require further scrutiny to clarify each model’s validity range. Real axons, for instance, display undulations and calibre variations [33], which can contribute to non-Gaussian diffusion. When such structural irregularities occur at length scales comparable to the diffusion length, the assumption of Gaussian diffusion within compartments may no longer hold. Their influence can be assessed by examining the time dependence of diffusivity [33, 34]. Additional microstructural features (such as myelin, vasculature, membrane permeability [35, 36], and the presence of glial cells [37, 38]) are also often neglected. For the diffusion times typically used in dMRI acquisitions (20–50 ms), the myelin sheath is generally assumed to act as an effective barrier that restricts exchange between the IAS and EAS, thereby minimising its contribution to the diffusion signal. However, unmyelinated axons, which permit water exchange, become increasingly common in pathological conditions such as multiple sclerosis [4], potentially introducing bias into SM parameter estimates that rely on the assumption of negligible permeability. Greater validation is therefore required to establish the range of conditions under which the SM can be reliably applied [28, 39].

Validating microstructural models by comparing their estimates with histology has been attempted in multiple studies, but this approach is far from straightforward. Indeed, histological tissue processing can unpredictably alter the underlying microstructure [40], introducing uncertainties that compromise the reliability of the chosen gold standard. An alternative strategy is to validate models by experimentally detecting specific functional forms in animal or human dMRI data (a paradigm borrowed from the physical sciences [13]) as was done for the stick model via its characteristic power law 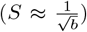 [41]. Synthetic signals generated directly from the analytical equations of the model of interest can also be compared with experimental data, but this approach cannot capture model failures when the true microstructure diverges from the model’s assumptions. Nonetheless, this technique was adopted for the validation of the Neurite EXchange Imaging (NEXI) model [42] for example. Furthermore, several studies have used physical phantoms (or hardware phantoms) [43–46], which incorporate an NMR-visible liquid alone or combined with NMR-invisible materials to produce well-defined microstructural properties. Physical phantoms come with several drawbacks, including complex chemical preparation, limited material availability, the need for specialised equipment, and a lack of standardisation [47]. In this work, we pursue yet another strategy, which is model validation using numerical simulations. This approach, increasingly popular in recent years, entails generating a numerical substrate (or virtual tissue) mimicking WM and then running Monte Carlo simulations to model water diffusion and synthesise dMRI signals. The advantage with this approach is that we may verify whether the estimated parameters are accurate relative to known ground truth (GT) features [47]. Despite notable progress in realistic substrate generation with various tools [34, 48–52], this technique has so far been applied mainly to simple substrates in which axons are represented as cylinders and lack realism [20, 43, 46, 53–61]. Conversely, some studies [33, 62, 63] have used numerical substrates obtained by segmenting electron-microscopy data (or confocal microscopy [64]), which provide highly realistic microstructural representations. However, this approach offers limited control over the substrate, since it is derived from real tissue, and it is considerably more expensive to acquire than generating numerical substrates “from scratch” with the tools mentioned above.

The aim of our study is thus to investigate the robustness and sensitivity of the SM parameter estimates in scenarios that deviate from the model assumptions using highly realistic numerical substrates. For this, we use the previously developed CATERPillar tool [34] for the generation of realistic numerical WM substrates and the Monte Carlo Diffusion and Collision (MC/DC) simulator [65]. We evaluate the performance of the SM in estimating parameters across substrates with different levels of complexity. To achieve this, separate analyses were performed on various types of substrates, each analysis representing a specific complexity level with a single varying parameter. These included variations in axonal undulation, beading, volume fraction, permeability, orientation dispersion, myelin volume fraction and the addition of permeable astrocytes. We hypothesise that incorporating complex axon morphologies, permeability, myelin and astrocytes will challenge the SM’s parameter estimation, as these factors are not accounted for in the model [30, 37, 38, 66].

## 2 Methods

### 2.1 Growing substrates with CATERPillar

Numerical substrates representing WM tissue were generated using our newly developed and publicly available CATERPillar framework [34]. These substrates could be populated with various cell types, including myelinated axons, non-myelinated axons, and glial cells (astrocytes, oligodendrocytes, etc.). To enhance biological realism, the cellular morphologies incorporated complex features such as axonal undulations and beading, parametrised to match histological measurements [34]. For this study, we simulated substrates with a volume of L = (150 *µ*m)^3^, using the default parameters summarised in Table 1.

**Table 1.**
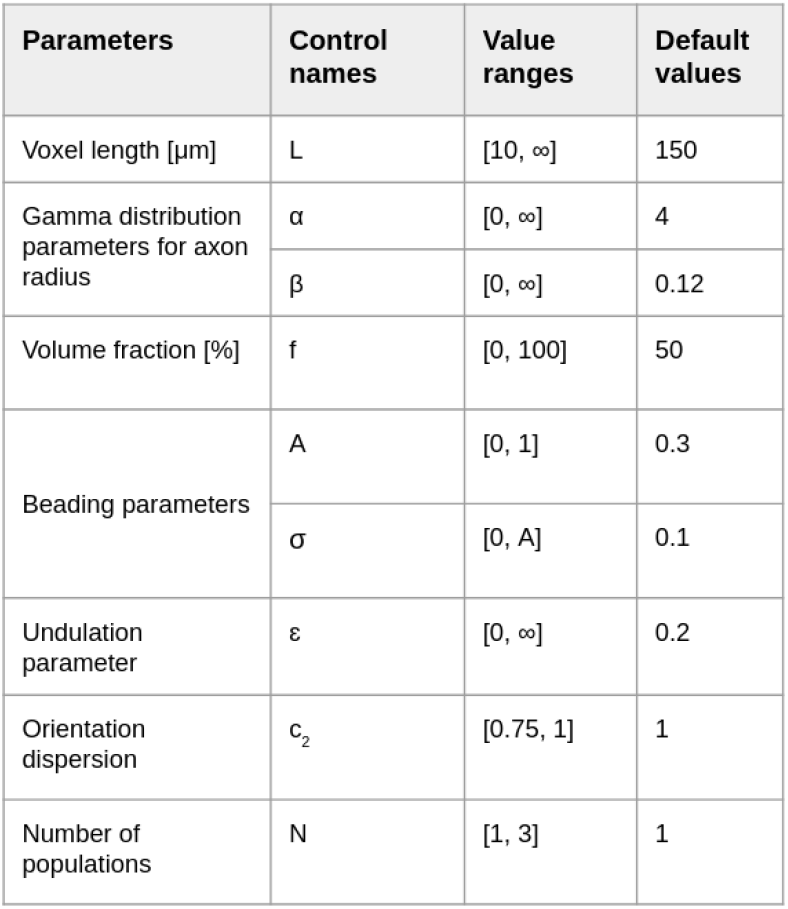
Default values for substrates generated by CATERPillar. These parameters lead to the creation of realistic axons as demonstrated in [34]. This Gamma Distribution with *α* = 4 and *β* = 0.25 yields a mean axonal radius of 1 *µm* and a mode of 0.75 *µm*. Setting A = 0.3 and *σ* = 0.1 for the beading parameters produce a coefficient of variation (CV) in axonal radius of approximately 0.2 along their length. The undulation parameter *ϵ* results in an undulation coefficient of 1.1. The orientation dispersion, here defined as *c*_2_ =*<* cos *ϕ*^2^ *>*, can be converted to *p*_2_ via the relationship *p*_2_ = 1.5*c*_2_ − 0.5.

The methods for growing numerical WM substrates with CATERPillar are described in detail in [34]. Briefly, axonal growth was initiated by assigning each axon a starting point on one plane and an attractor on the opposite plane with a pre-defined volume fraction *f* . Each axons mean radius (*R*_*mean*_) was sampled from a Gamma Distribution with controllable parameters (by default *α* = 4 and *β* = 0.12). During growth, spheres were sequentially added to form a chain of overlapping elements, extending towards the attractor. Undulations were introduced by setting a non-zero value for *ϵ*, a parameter that modulates the strength of attraction towards the target. Increasing *ϵ* weakens this attraction, leading to more pronounced undulations along the axon’s trajectory, as shown in Figure 2. For instance, with *ϵ* = 0.2, the typical undulation coefficient (or tortuosity defined as the ratio between the total axon length and the straight-line distance between its endpoints) is approximately 1.1. Beading was implemented in a stochastic manner by sampling the radius of each subsequent sphere from a Gaussian distribution at every growth step. The distribution was centred on the radius of the preceding sphere and constrained to lie within *R*_*mean*_ ± *A* × *R*_*mean*_. The parameter *A* thus set an upper bound to the amplitude of the beading relative to *R*_mean_. On the other hand, *σ* controlled the spatial frequency of the beading, as the standard deviation of the Gaussian distributions were equal to *σ* × *R*_*mean*_. The *σ* parameter was fixed in the previous version of CATERPillar [34], but is now a tunable user-defined parameter. Setting A = 0.3 and *σ* = 0.1 produced a coefficient of variation (CV) in axonal radius of approximately 0.2 along their length, consistent with measurements from electron microscopy [67, 68]. These values were therefore adopted as the default parameters, inducing beading as illustrated in Figure 1.

**Figure 1.**
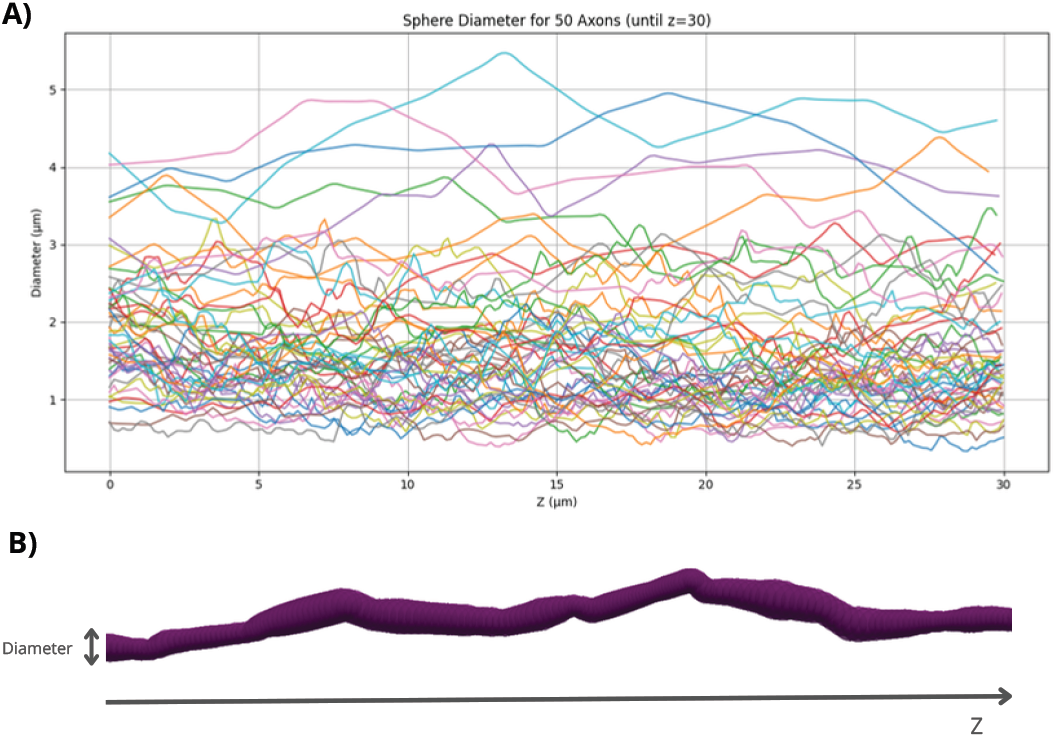
A) The axonal diameter variation displayed along the main direction (z) for 50 random axons. B) Illustration of a single axon with varying diameter along its length.

**Figure 2.**
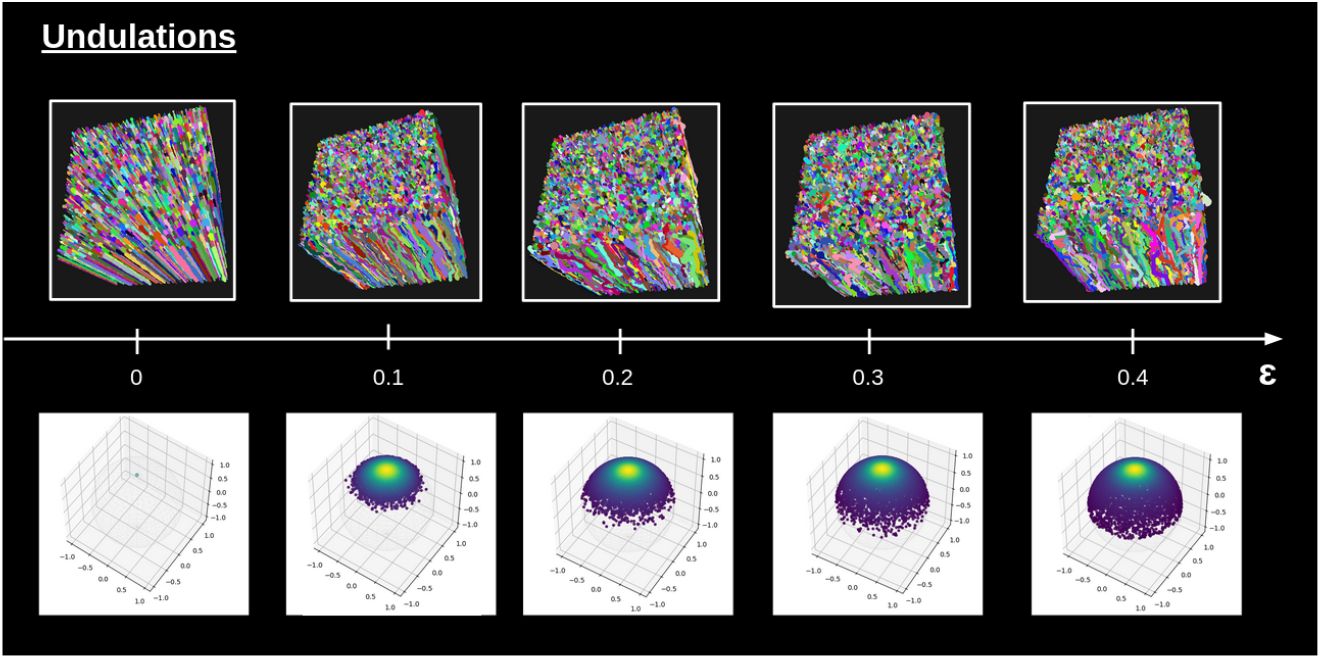
The axonal undulations are governed by the parameter *ϵ*. At each growth step, the angles determining the position of the next sphere are sampled from a Gaussian distribution centred on the axon’s attractor direction. The value of *ϵ* sets the standard deviation of this distribution. When *ϵ* approaches zero, axons extend almost perfectly straight towards their attractor, whereas larger values of *ϵ* introduce increasing deviations, resulting in more pronounced undulations along the axon’s trajectory. The vectors corresponding to the sampled angles are displayed on the spherical surfaces in the last row. A larger value of *ϵ* permits greater deviation of the vectors from the attractor located at the pole.

Axonal fanning was also simulated using a Watson distribution parameterised by *c*_2_. In this model, each axon’s attractor was oriented at a specified angle relative to its origin, allowing controlled degrees of fanning. The *c*_2_ parameter could be converted to the SM orientation dispersion index *p*_2_ through the relationship *p*_2_ = 1.5 *c*_2_ − 0.5. To generate greater overall dispersion, up to three perpendicular fibre populations (*N* = 3) could be created.

After axonal growth was completed, myelin could be introduced by generating inner spheres within the pre-existing ones. The space between the inner and outer spheres represented the myelin sheath and was subsequently excluded from water molecule placement during the Monte Carlo simulation.

As for glial cells, the somas were placed prior to the axonal growth, while the glial processes (or branches) were grown from the soma towards an assigned attractor after the axons were placed. The mean/standard deviation for the soma radius, number of primary processes, mean/standard deviation for the process length and the volume fractions for the soma and processes can be set by the user, so that the morphological features of the glial cells can match any glial cell of interest. In a part of this work, we populated our substrates with astrocytes, so adapted the morphological parameters so that they matched histological findings, as also previously done in [34]. Example of astrocytes present in our substrates are shown in Figure 3.

**Figure 3.**
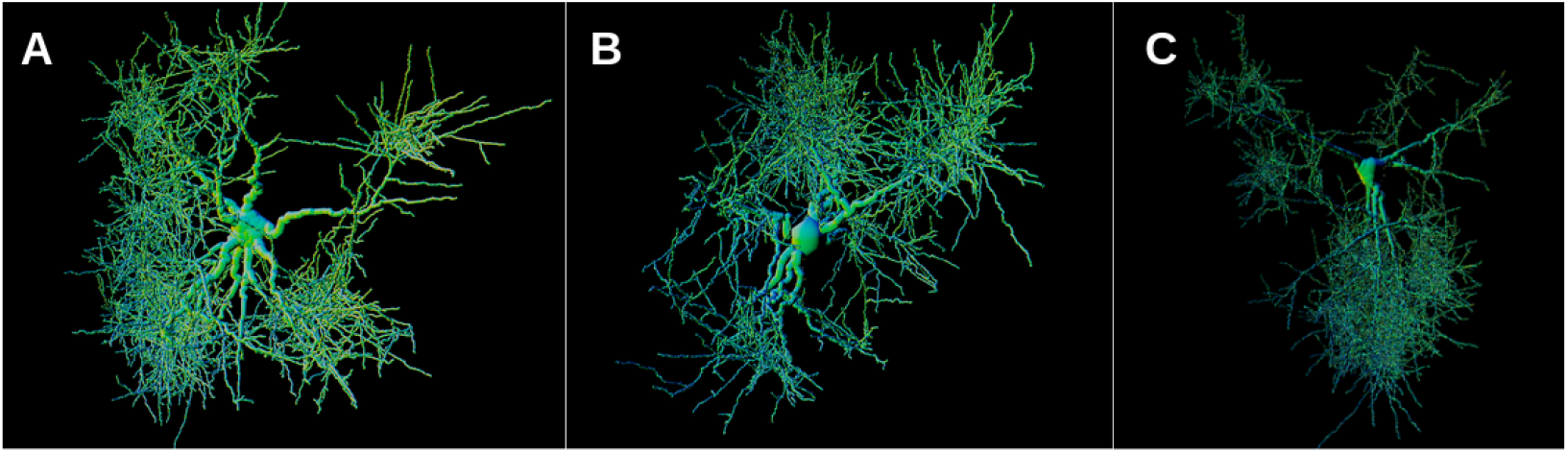
Example of astrocytes generated with CATERPillar.

### 2.2 Monte Carlo Simulations

Monte Carlo simulations were conducted within the generated voxels with the MC/DC tool [65]. This simulator tool produced diffusion signals from a numerical substrate, neglecting *T*_2_ effects and myelin water, and therefore did not place water particles within the myelin compartment. Simulations used bulk diffusivities of 2.5 *µ*m^2^*/*ms for IAS and 1.5 *µ*m^2^*/*ms for EAS. These diffusivity values were chosen to align with previous studies reporting IAS diffusivities near 2.2 *µ*m^2^*/*ms [69–71] and supporting the condition *D*_*a*_ *> D*_*e*∥_ [28, 70, 72]. Random walkers were initialised in a central (110 *µ*m)^3^ region within the (150 *µ*m)^3^ cube to avoid edge effects. A step size of 0.1 *µ*m (3 times smaller than the smallest sphere diameter) yielded 38,500 and 23,100 steps for IAS and EAS, respectively. Simulations used a walker density of 1 walker */µm*^3^, which lead to a total of 1,330,000 walkers when considering (110 *µ*m)^3^. This density of walkers was proven to generate stable signals in Figure **??**.

Once the trajectories were obtained, we generated synthetic dMRI signals with a pulse-gradient spin-echo (PGSE) sequence [73]. A wide pulse was utilised to represent clinical scanners, employing Δ = 55.5 ms and *δ* = 16.5 ms. Sixty unique gradient directions per shell were generated with optimal angular coverage using the method outlined by [74]. The b-values utilised were 0, 500, 1000, 2000, 3000 and 5000 *s/mm*^2^ for SM estimations. Rician noise was added to the signal with signal-to-noise ratios (SNRs) of the non diffusion-weighted signal (b=0) of 50 and 100, with 1000 random noise realisations per noiseless signal.

### 2.3 Estimating SM parameters

The objective of this analysis is to evaluate how morphological features that are accounted or unaccounted for explicitly in the model influence the SM estimations. When estimating the SM parameters, we did not consider the free water (FW) compartment, for which multi-echo time data was needed to solve model degeneracy [5]. By fitting the two-compartment SM to dMRI data, several microstructural parameters were estimated:

1. Axonal Water Fraction (*f*): Represents the relative fraction of MRI signal originating from the IAS. This fraction excludes any myelin water contribution, as myelin water has very short *T*_2_ and is MR-invisible in dMRI due to the typically long echo time used [4, 75, 76]. When the SM is applied to real dMRI data, the relative signal fraction is weighted by the *T*_2_ of each compartment. In our simulations however, we did not place particles inside the myelin layer, and we neglect *T*_2_ relaxation. Caution should be taken when interpreting *f*, as the signal fraction is not necessarily equal to the axonal volume fraction (especially in the presence of myelin or compartment-specific *T*_2_ relaxation times).
2. Intra- and Extra-Axonal Diffusivities (*D*_*a*_, *D*_*e*∥_, *D*_*e*⊥_): These parameters quantify the diffusion coefficients within both the intra- and extra-axonal spaces. Diffusivity can be measured either parallel (∥) or perpendicular (⊥) to the primary axonal orientation.
3. Fibre Orientation Distribution Function (fODF): Captures the dispersion of axonal orientations within a voxel. fODF anisotropy is minimally characterised by the *l* = 2 order rotational invariant *p*_2_.

The TMI Python toolbox (https://nyu-diffusionmri.github.io/DESIGNER-v2/docs/TMI/SMI/) [5, 12, 77] was used to apply the SM to synthetic dMRI signals. As described in [5], the rotational invariants are first computed using a least-squares estimator, then the parameters are obtained using a data-driven machine learning regression. Specifically, a cubic polynomial regression model is employed [14], as it has been demonstrated to be more effective than neural networks for this application.

Our analysis can be divided into several parts:

1. **Axonal Morphology:** Here, the aim was to assess how morphological features such as undulations and beading impact the SM estimations. The analysis comprised two parts: variation of undulation strength and variation of beading strength. The intra-axonal volume fraction *f* was kept constant at 0.5 in both parts.
  (a) Undulations (or tortuosity) were introduced by setting the *ϵ* parameter in CATERPillar to values above zero, ranging from 0.1 to 0.4 across four different substrates. This resulted in undulation coefficients varying between 1.05 and 1.3, which are consistent with values reported for WM tissue [78, 79]. All other parameters were kept to their default values (Table 1).
  (b) Axonal beading (or calibre variation) was introduced by varying the *σ* parameter in the CATERPillar tool. This parameter ranged from 0 to 0.4 in four different substrates, with higher values producing greater beading strengths. The degree of beading was quantified by the mean CV of the axonal radius along each axon’s length, averaged across all axons within the voxel, which ranged between 0.1 and 0.4. Comparable CV values have been reported in previous electron microscopy studies of WM tissue [67, 68], with typical mean CVs around 0.2-0.3.
2. **Axonal Packing:** The axonal volume fraction *f* varied from 0.1 to 0.6 in 6 different substrates.
3. **Orientation Dispersion:** Substrates with a constant *f* of 0.3 were generated while systematically varying the degree of fibre dispersion. All other parameters were assigned their default values (Table 1). In total, five substrate configurations were simulated: (A) three mutually perpendicular fibre populations, (B) two perpendicular populations arranged as planar sheets, (C) two perpendicular and intersecting populations, (D) a single fibre population exhibiting fanning modelled using a Watson distribution, and (E) a single population without fanning. The highest degree of orientation dispersion (i.e., the lowest *p*_2_) was observed in the configuration with three perpendicular populations (A), as this arrangement produced an approximately isotropic voxel.
4. **Cell Permeability:** In this part, we aimed to assess the impact of cell permeability on the SM results. Permeability was applied to axons and to astrocytes in two separate analyses.
  (a) We used a substrate with undulating and beaded axons with a packing *f* = 0.5. The permeability of these axons ranged from 0 (no permeability) to 4 × 10^−5^*m/s*, as previously reported in experimental studies [80].
  In a substrate containing an axonal packing of 0.5, we this time also included astrocytes. These cells are composed of a soma (representing 1% of the total voxel volume) and of processes (representing 6% of the total voxel volume). The volume fractions for astrocytes were chosen to match those reported in [81] for deep WM. The astrocyte permeability ranged from 0 to 10^−4^*m/s* to also match results obtained from [80].
5. **Myelin:** A substrate was generated consisting of axons surrounded by an outer layer representing the myelin sheath. The intra-axonal volume fraction (AVF) combined with the myelin volume fraction (MVF) was approximately 0.5, with 0.24 corresponding to AVF and 0.26 to the MVF. The myelin thickness for each axon was determined as a function of the axonal radius, according to the relationship described by [67]. Axonal radii were sampled from a Gamma distribution with parameters *α* = 2 and *β* = 0.12, which was different from the default values of *α* and *β* so that we this time mimic the distribution of the inner rather than outer-axonal radii. This substrate composed of realistically myelinated axons served as the baseline model. Subsequently, the myelin thickness of all axons was progressively reduced by − 25%, − 50%, and − 75%, as well as a final case with no remaining myelin. These modifications yielded MVFs of 0.263, 0.225, 0.125, 0.075 and 0 respectively. It is important to distinguish AVF, AVF+MVF and the axonal water fraction *f* estimated by the SM. Since myelin is considered a compartment MR-invisible, it does not contribute to the total dMRI signal. Consequently, when myelin is present in the substrate, the following relationships hold:

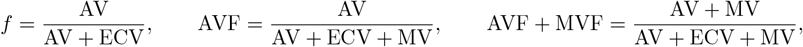

and therefore, if MVF ≠ 0, then *f* ≠ AVF ≠ AVF + MVF, where AV is the intra-axonal volume, ECV the extra-cellular volume and MV the myelin volume.
6. **Diffusion Time:** In this part, a single substrate with *f* = 0.5 was used, and the simulation was performed once. PGSE sequences with identical *δ* but varying Δ values (20, 35, 50, 75, 100, and 125 ms) were used to sample signals at different diffusion times. Each PGSE sequence was applied to the same initial trajectory, which was then cropped according to the sequence’s duration. The signal acquired at Δ = 125 ms served as the reference for estimating *D*_∞_ across all diffusivity measurements.

The SM estimates obtained in each of these configurations were compared to their corresponding effective GT values. These values for *f* and *p*_2_ were computed directly from the substrate geometry, in case the they deviated from the target values. Since all substrates exhibited some degree of axonal undulation, the effective GT *p*_2_ values were consistently below 1, which corresponds to perfectly aligned cylindrical fibres. The GT diffusivity metrics *D*_*e*⊥_, *D*_*e*∥_, and *D*_*a*_ were derived from the simulated water molecule trajectories generated by the Monte Carlo simulator. In the case of permeable cells, the water molecules from different compartments could not be separated, so the GT compartment specific diffusivity could not be obtained. In these cases, we instead computed *D*_⊥_ and *D*_∥_ which indicated the perpendicular and parallel diffusivity of both compartments combined.

## 3 Results

The following subsections present the effects of specific parameters on the SM parameter estimates. Each parameter was varied independently while all others were kept constant. Figures 5–12 illustrate the influence of each parameter on the SM estimates, both with and without the addition of noise. Figure 12 summarises the overall impact of all parameters on the SM estimates (A) and the error relative to their corresponding GT (B) in the absence of noise.

### 3.1 Axonal undulations

As previously described, the axonal undulation within a voxel was quantified as the ratio between the axonal length and the straight-line distance between its endpoints. The results shown in Figure 4 indicate that the SM estimates of *p*_2_, *D*_*a*_, and *D*_*e*∥_ decreased by 15%, 10%, and 11%, respectively, with increasing undulation coefficients, whereas *D*_*e*⊥_ increased by 11%. In contrast, the estimated *f* varied by only 1.1%, remaining highly stable. The relative error of the estimates with respect to the effective GT were consistent across all levels of undulation and varied by less than 2%, except for *D*_*a*_, whose overestimation increased from 6.4% to 13% at the maximum undulation strength.

**Figure 4.**
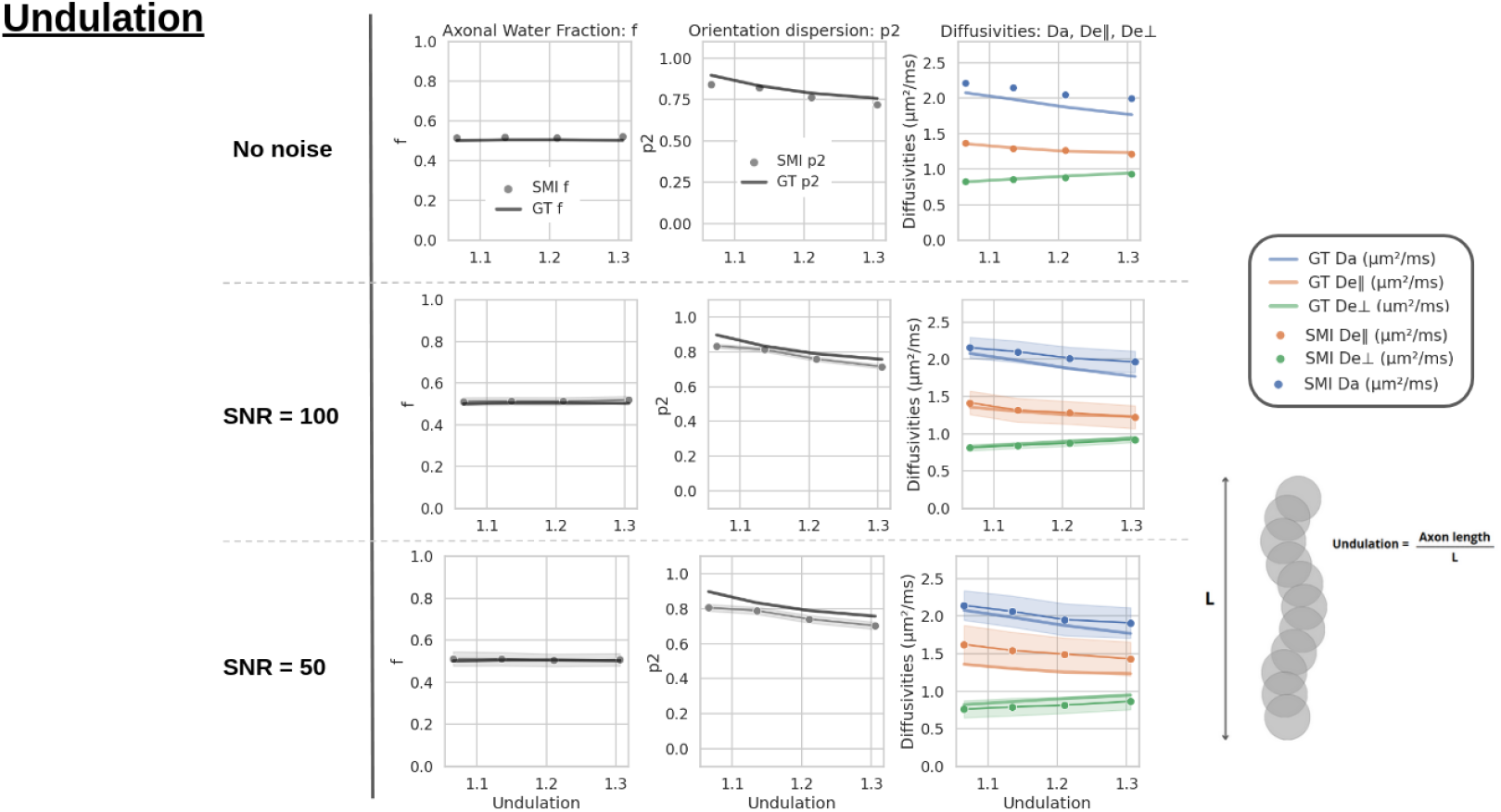
SM estimates of *f, p*_2_, *D*_*e*∥_, *D*_*e*⊥_, and *D*_*a*_ are shown together with their corresponding GT values as a function of the mean undulation coefficient for each voxel. When noise is included, the shaded areas represent the standard deviation across 1000 repetitions.

### 3.2 Axonal beading

Beading was modulated across subjects by varying the parameter *σ* (Figure 1) from 0 to 0.3. The degree of beading was quantified as the mean CV of the axonal radius along each axon’s length, averaged across all axons within the voxel. As shown in Figure 4, increasing mean CV from 0 to 0.3 led to rises in the estimated *f* and *D*_*e*⊥_ by 20% and 5.9%, respectively, while *p*_2_, *D*_*a*_, and *D*_*e*∥_ decreased by 26%, 19%, and 3%. The relative errors of *f, p*_2_, and *D*_*a*_ with respect to their GT were more strongly influenced by beading (with variations over 9%) than those of *D*_*e*⊥_ and *D*_*e*∥_, which varied by less than 2%. Specifically, *f* and *D*_*a*_ were increasingly overestimated with higher beading amplitudes, whereas *p*_2_ became progressively more underestimated.

### 3.3 Axonal packing density

In Figure 6, the SM estimates varied with increasing GT *f* . Excellent accuracy and precision were observed between estimated *f* and its GT, which explains its increase by 82% over the entire range. The estimates of *p*_2_ displayed a bell-shaped evolution, and showed a maximum relative error of −30% due to underestimation at the lowest packing density. *D*_*a*_ also displayed a mild bell-shaped pattern with increasing GT *f*, which became monotonous in the presence of noise. *D*_*a*_ was typically overestimated with respect to its effective GT for lower axonal packing densities. As expected, all EAS diffusivity estimates decreased with packing, by 17 % for *D*_*e*∥_ and 45% for *D*_*e*⊥_.

**Figure 5.**
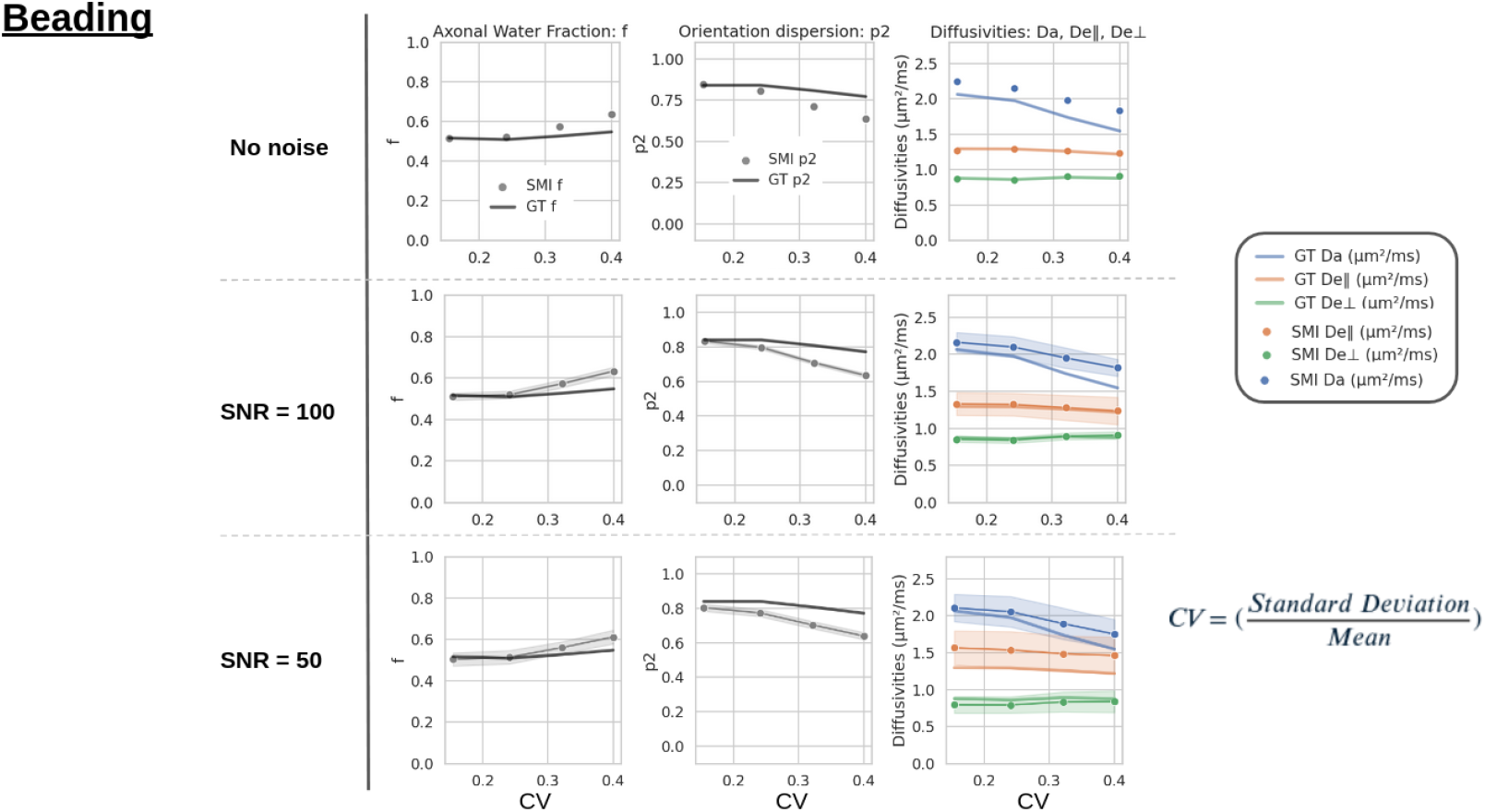
SM estimates of *f, p*_2_, *D*_*e*∥_, *D*_*e*⊥_, and *D*_*a*_ are shown together with their corresponding GT values as a function of the mean CV for each voxel. This CV quantified the beading strength of the axons, which was modulated across subjects by varying the parameter *σ* from 0 to 0.3. The mean CV of the axonal radius was obtained by averaging the CV for all axons within the same voxel. When noise is included, the shaded areas represent the standard deviation across 1000 repetitions.

**Figure 6.**
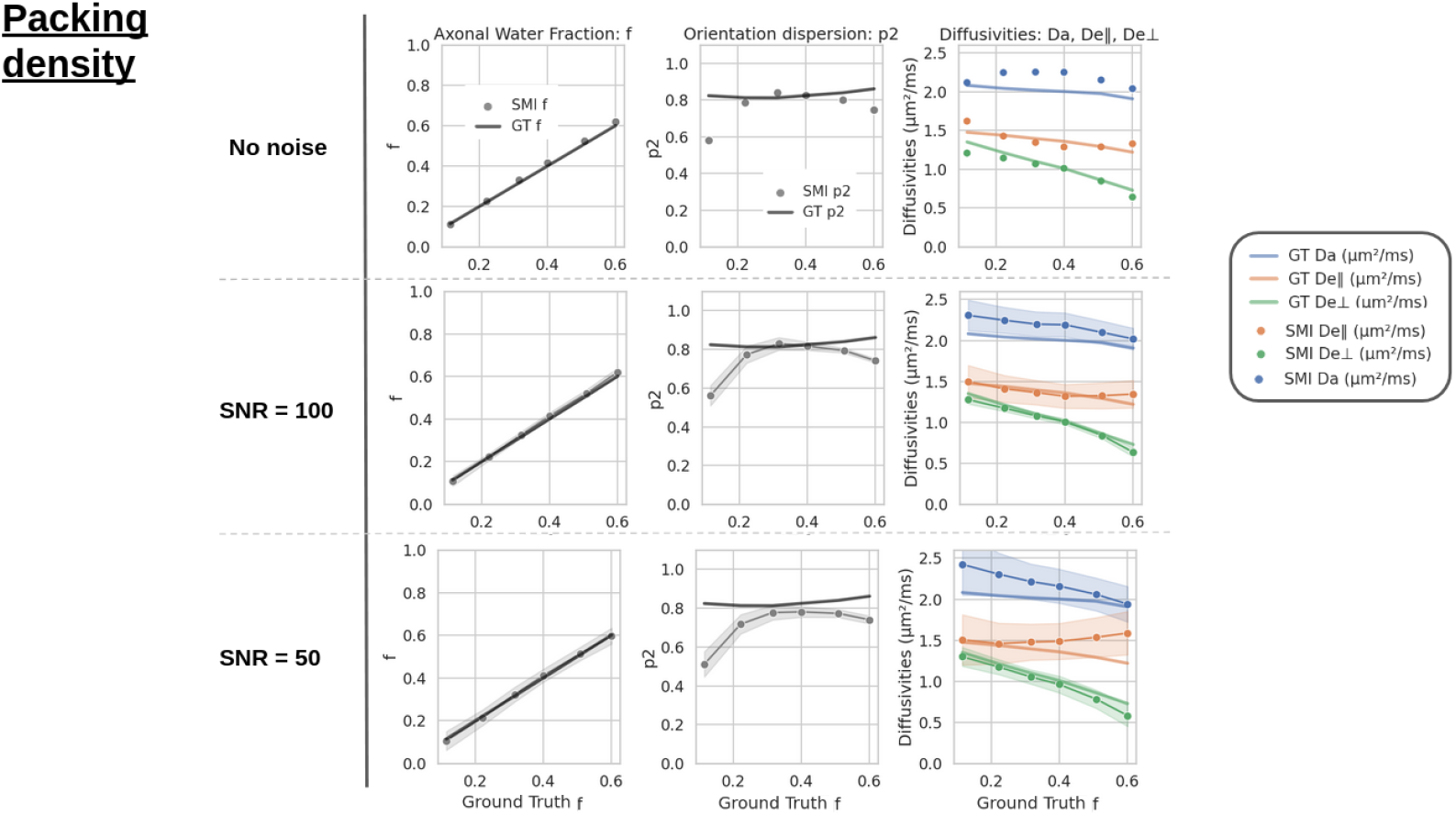
SM estimates of *f, p*_2_, *D*_*e*∥_, *D*_*e*⊥_, and *D*_*a*_ are shown together with their corresponding GT values as a function of the GT axonal volume fraction *f* . When noise is included, the shaded areas represent the standard deviation across 1000 repetitions.

### 3.4 Orientation Dispersion

Orientation dispersion, represented by the GT *p*_2_, was varied by including different numbers of fibre populations and by inducing axonal fanning. Substrate (A) contained three perpendicular fibre populations, while substrates (B) and (C) each included two perpendicular populations arranged in sheet-like and interwoven configurations, respectively. Substrate (D) consisted of a single fibre population with fanning, modelled using a Watson distribution, and substrate (E) featured a single population with no fanning. As shown in Figure 7, B and C revealed very similar parameter estimates, as they were both composed of two perpendicular fibre populations. This similarity suggests that the SM is not sensitive to different crossing configurations.

**Figure 7.**
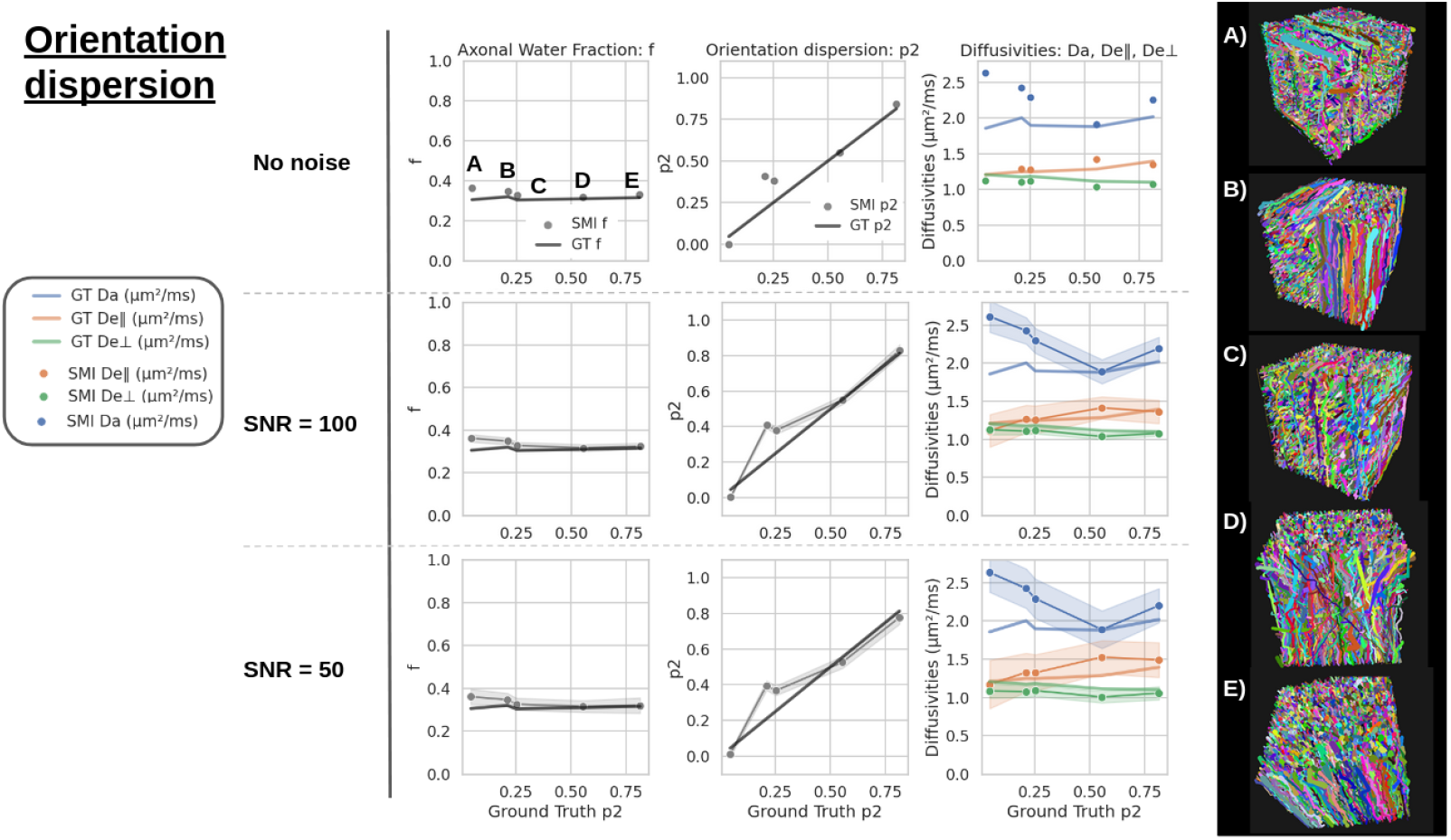
SM estimates of *f, p*_2_, *D*_*e*∥_, *D*_*e*⊥_, and *D*_*a*_ are shown together with their corresponding GT values as a function of the mean GT *p*_2_ for each voxel. Substrate (A) contained three perpendicular fibre populations, while substrates (B) and (C) each included two perpendicular populations arranged in sheet-like and interwoven configurations, respectively. Substrate (D) consisted of a single fibre population with fanning, modelled using a Watson distribution, and substrate (E) featured a single population with no fanning. When noise is included, the shaded areas represent the standard deviation across 1000 repetitions.

The estimated *p*_2_ closely matched its GT, except for the two-fibre configurations (B and C) (*p*_2_ ≈ 0.25), where it was overestimated. With increasing *p*_2_, corresponding to greater axonal alignment, the estimated *f* and *D*_*e*⊥_ decreased by 8.8% and 6.1%, respectively, while *D*_*e*∥_ increased by 15% across the full range, staying broadly in line with their effective GT values. Indeed, the relative errors of *D*_*e*⊥_ and *D*_*e*∥_ remained low and stable, with variations not exceeding 5%. At low GT *p*_2_ values (A–C) however, *D*_*a*_ was overestimated, with maximum relative errors of 42%, yielding a fluctuation of 17% across the *p*_2_ range that was not matched by the effective GT values estimated from the trajectories.

### 3.5 Axonal Permeability

In Figure 8, the permeability of all axons was increased from 0 (impermeable) to 4 × 10^−5^ m*/*s in steps of 1 × 10^−5^ m*/*s. This led to a 75% decrease in the estimated *f*, despite its GT remaining constant, resulting in a maximum relative error of 75% when the permeability equals to 4 × 10^−5^ m*/*s. In contrast, *p*_2_ and *D*_*a*_ remained relatively stable, increasing by only 5.5% and 4.6%, respectively. The extracellular diffusivities *D*_*e*∥_ and *D*_*e*⊥_ varied more substantially, with *D*_*e*∥_ increasing by 26% and *D*_*e*⊥_ decreasing by 39%. A direct comparison between the diffusion estimates and their GTs was not possible, as the presence of permeability prevented clear separation of intracellular and extracellular trajectories. Instead, the GT *D*_⊥_ and *D*_∥_, which are not compartment-specific, were reported and both increased slightly with axonal permeability. At zero permeability, *D*_⊥_ was positioned between *D*_*e*⊥_ and *D*_*a*⊥_ = 0, while *D*_∥_ was positioned between *D*_*e*∥_ and *D*_*a*_. In the noiseless case, *D*_*e*∥_ and *D*_*e*⊥_ converged towards their average counterparts (*D*_∥_ and *D*_⊥_), while *D*_*a*_ slightly increased but contributed gradually less to *D*_∥_ due to underestimation of a restricted stick compartment (*f*).

**Figure 8.**
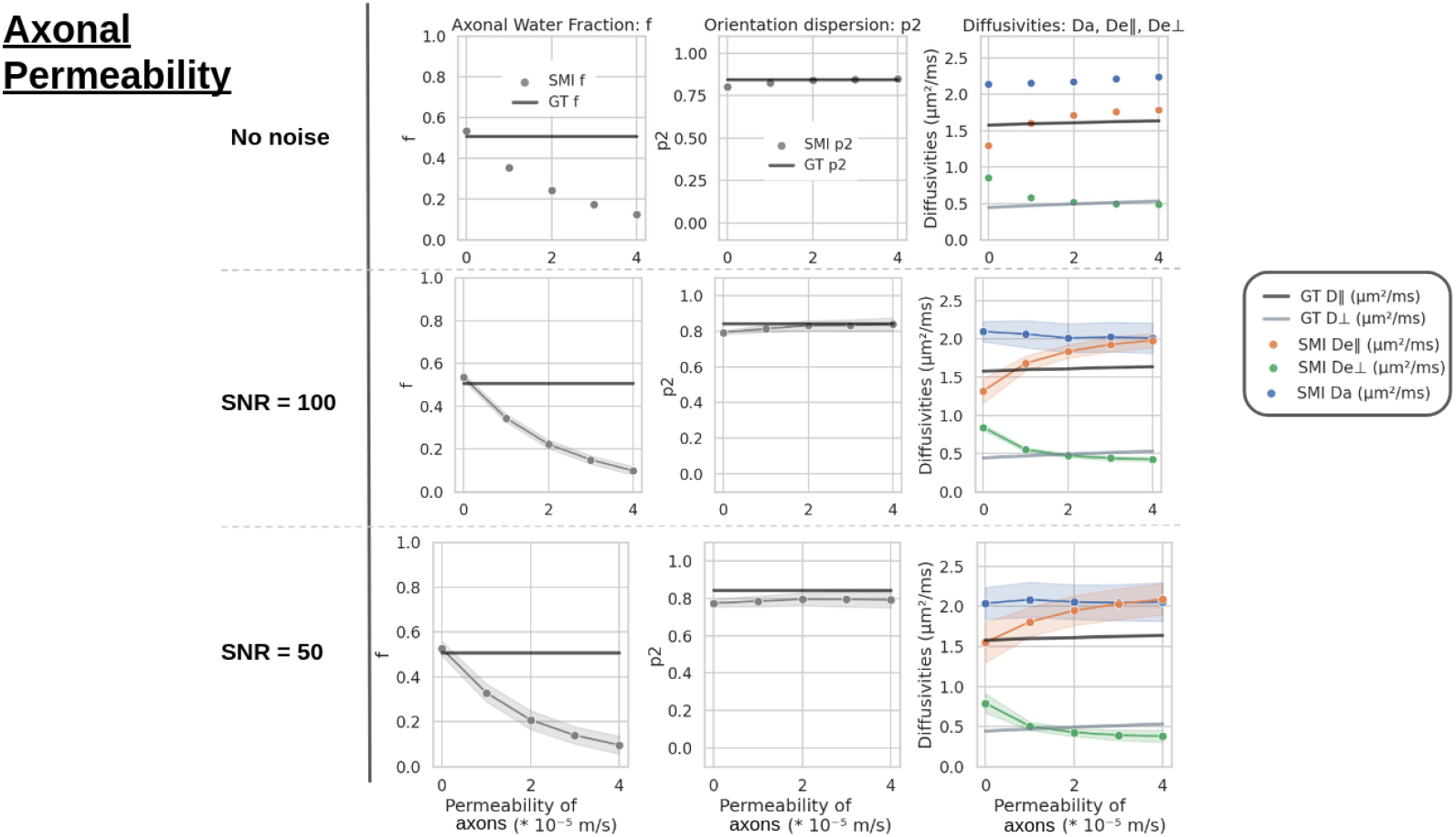
SM estimates of *f, p*_2_, *D*_*e*∥_, *D*_*e*⊥_, and *D*_*a*_ are shown together with their corresponding GT values as a function of the permeability of the axons for each voxel. When noise is included, the shaded areas represent the standard deviation across 1000 repetitions.

### 3.6 Astrocyte Permeability

Astrocytes were included in voxels containing axons, with volume fractions and morphological parameters adjusted to match realistic astrocyte properties in deep WM, as described previously [81]. The permeability of astrocytes was varied from 0 (impermeable) to 10^−4^ m*/*s. In Figure 9, AVF denotes the axonal volume fraction, while AVF + ASVF represents the combined volume fraction of axons and astrocytes (processes and soma). Astrocytes had the greatest influence on the SM estimates at low permeability. Increasing astrocyte permeability improved the axonal estimation accuracy of both *f* and *p*_2_, as the relative error went from 35% to 9.8% for *f* and from -12% to 3.7% for *p*_2_. At low permeability values, *f* and *p*_2_ were overestimated and underestimated, respectively. The diffusivity estimates were largely stable, showing minor increases with astrocyte permeability with less than 3% for *D*_*a*_ and *D*_*e*⊥_, and a moderate increase of 4.7% for *D*_*e*∥_. *D*_∥_ and *D*_⊥_, which are not specific to a compartment, both increased by approximately 0.03 *µm*^2^*/ms* across the full range of astrocyte permeability.

**Figure 9.**
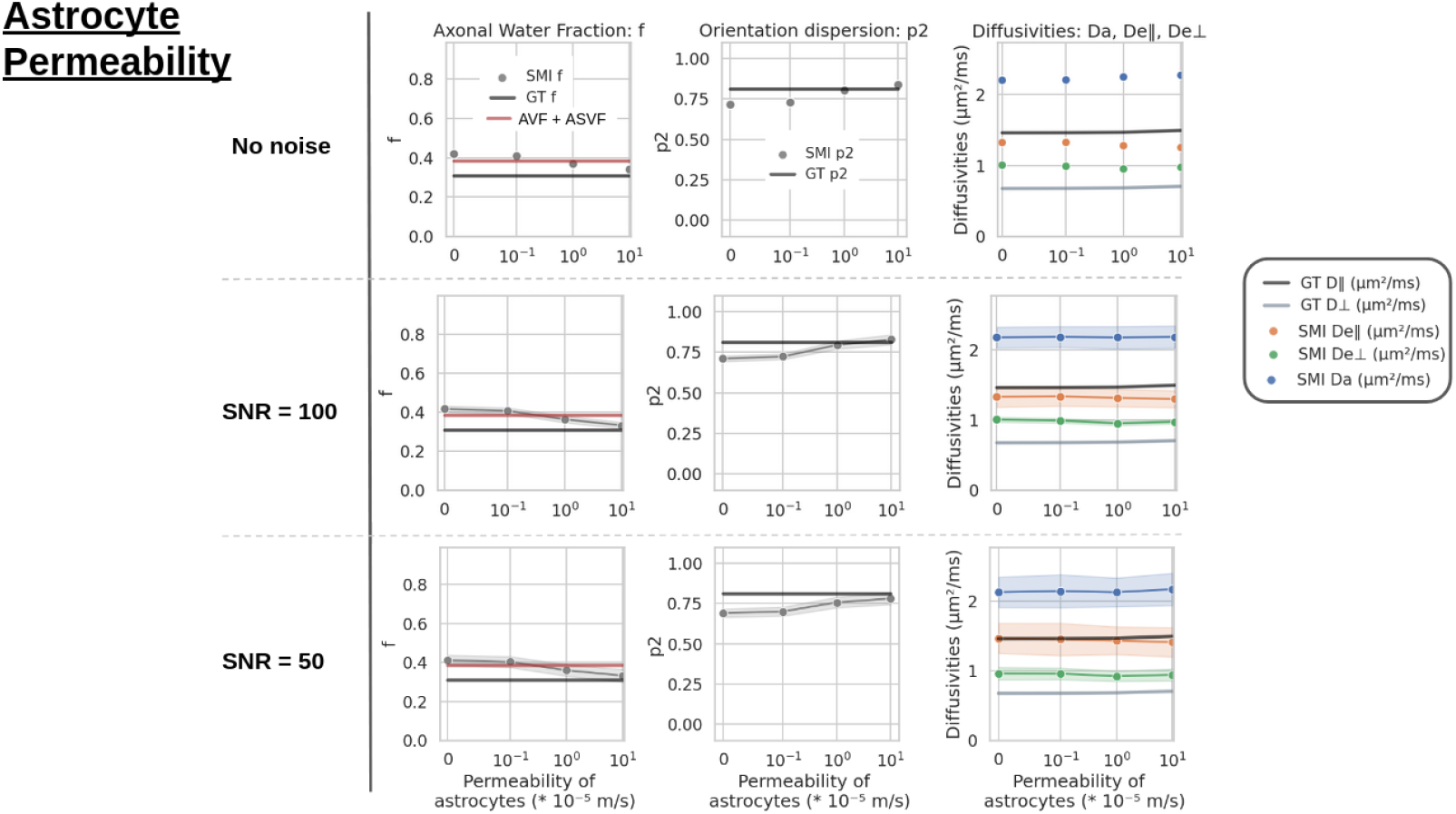
SM estimates of *f, p*_2_, *D*_*e*∥_, *D*_*e*⊥_, and *D*_*a*_ are shown together with their corresponding GT values as a function of the permeability of the astrocytes for each voxel. The astrocytes’ soma and processes took 1% and 6% of the total voxel volume. Additionally to GT *f*, the SM *f* can be compared to AVF + ASVF, which accounts for the volume occupied by both axons and astrocytes. When noise is included, the shaded areas represent the standard deviation across 1000 repetitions.

### 3.7 Myelin Volume Fraction

Figure 10 illustrates the behaviour of the estimates when a myelin sheath was introduced and progressively thickened until reaching a realistic myelin-to-axon radius ratio, as described by [67]. The estimated *f* increased by 39% across the full range and was positioned midway between AVF+MVF and AVF. It was overestimated relative to its GT, with a 13% error at the highest MVF. The parameter *p*_2_ was consistently underestimated but remained stable across varying MVF values (variation below 0.1%). In contrast, *D*_*a*_ was systematically overestimated and increased slightly by 1.7% despite a constant GT. *D*_*e*∥_ decreased slightly (by 4.4%) while *D*_*e*⊥_ decreased dramatically (by 33%) with increasing MVF. Their relative errors remained low.

**Figure 10.**
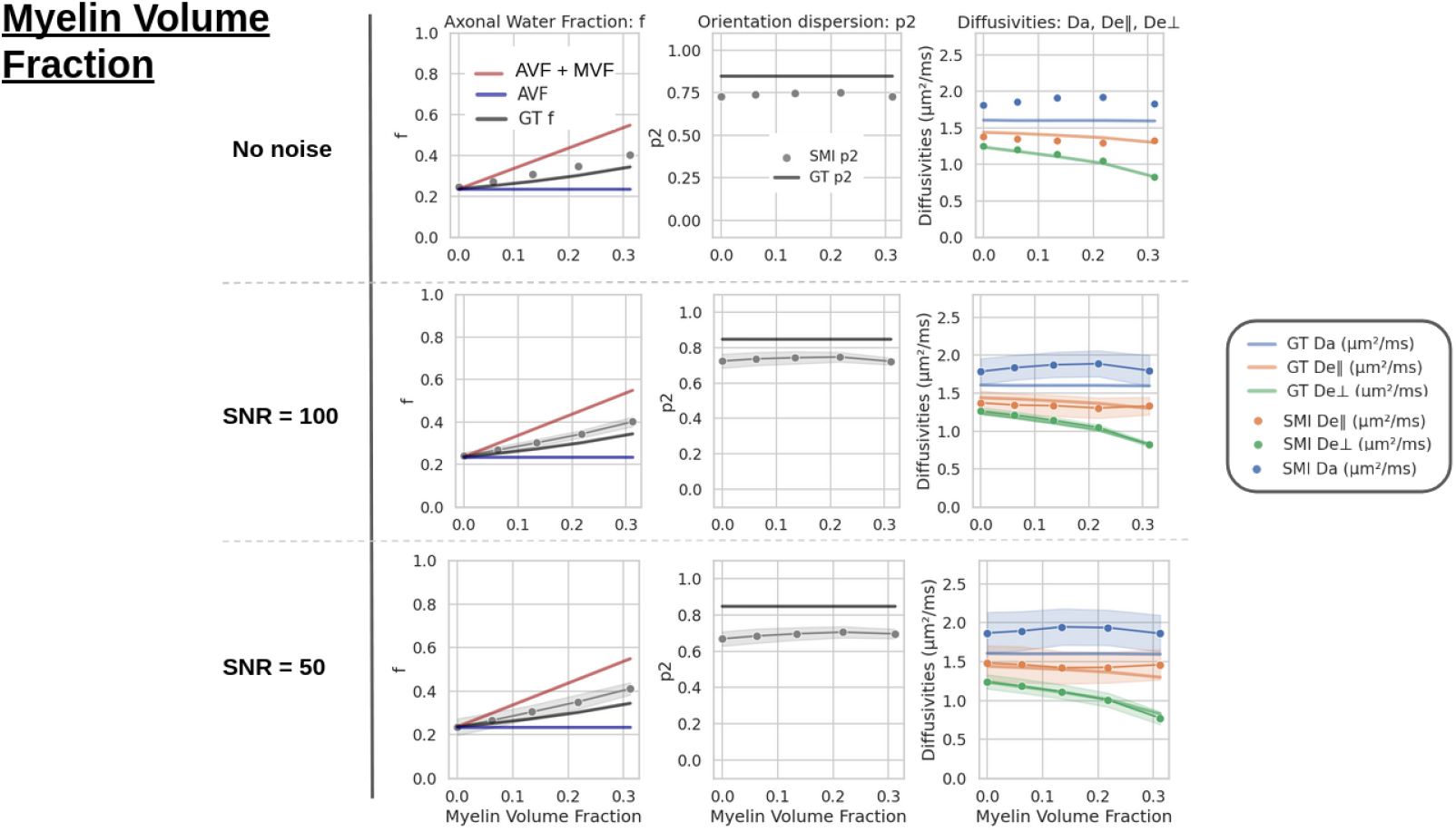
SM estimates of *f, p*_2_, *D*_*e*∥_, *D*_*e*⊥_, and *D*_*a*_ are shown together with their corresponding GT values as a function of the MVF for each voxel. GT f takes into account the reduced water volume with increasing MVF, thus increases despite a constant AVF. The AVF + MVF represents the real volume fraction of axons with myelin included. When noise is included, the shaded areas represent the standard deviation across 1000 repetitions.

### 3.8 Diffusion Time

The same substrate containing undulating and beading axons was used, with a packing density of 0.5. In this case, the substrate geometry remained fixed, and the diffusion time of the water particles was varied instead. Figure 11 shows the estimated parameters compared to their GTs, with the diffusivity GT computed at a longer diffusion time (0.12 s) and considered as *D*_∞_. All estimated parameters converged towards their GTs with increasing diffusion time. This convergence was achieved through a gradual decrease in the estimated values for all parameters except *p*_2_, which increased towards its GT. While *D*_*e*⊥_ was already in its long time limit even at short diffusion times, the axial diffusivities *D*_*a*_ and *D*_*e*∥_ showed measurable time-dependence, particularly up to Δ = 50 ms.

**Figure 11.**
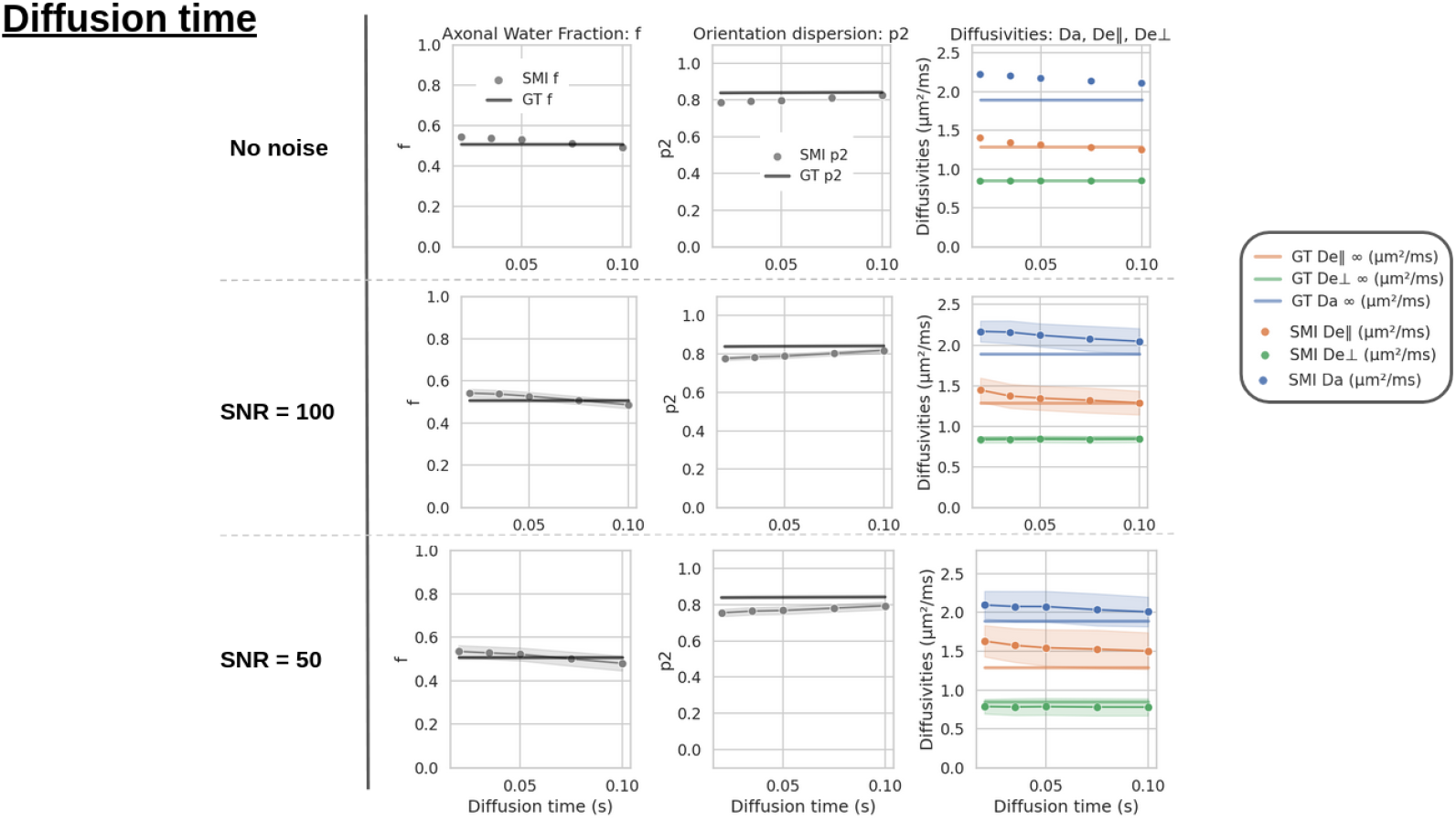
SM estimates of *f, p*_2_, *D*_∥_, *D*_*e*⊥_, and *D*_*a*_ are shown together with their corresponding GT values as a function of the diffusion time for each voxel. When noise is included, the shaded areas represent the standard deviation across 1000 repetitions.

**Figure 12.**
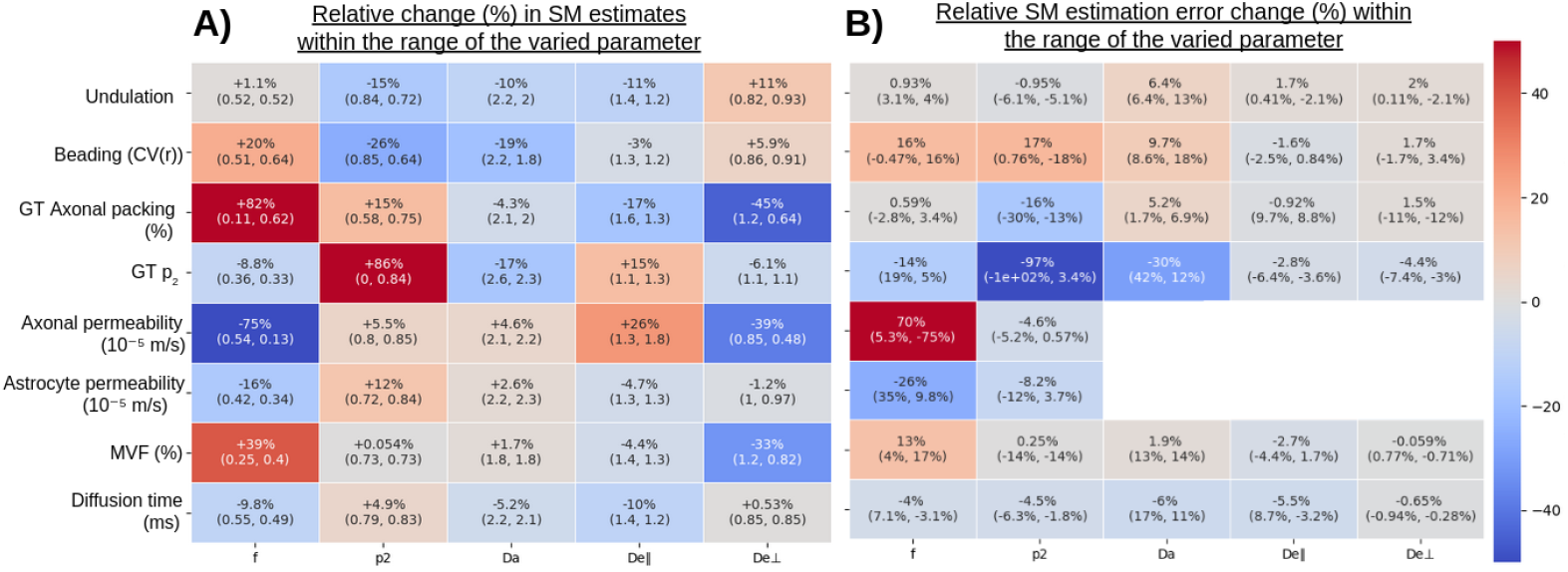
(A) Relative sensitivity of each SM estimate across the range of the varied parameter. The varied parameter was normalized, and each SM estimate was divided by its maximum value within this range. A linear regression was then fitted, and the resulting slope was expressed as a percentage. (B) Relative change in the error of each SM estimate across the range of the varied parameter. The error was defined as 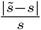, where 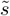 denotes the SM estimate and *s* the corresponding GT. The error was computed for the lowest and highest values of the varied parameter (shown in parentheses), and their difference is reported. This difference is negative when the variation decreases the SM error (i.e., improves accuracy) and positive when it increases the error. A value close to zero indicates that the parameter had little influence on the SM estimation error. Both (A) and (B) correspond to noiseless data.

### 3.9 Noise

In Figures 5–12, the addition of noise produced similar effects across most simulations. As the SNR decreased (i.e., as noise levels increased), the standard deviation widened accordingly. In addition, noise introduced a systematic bias as an underestimation of *p*_2_ and an overestimation of *D*_*e*∥_. The latter effect was particularly pronounced at an SNR of 50.

## 4 Discussion

In this work, we evaluated the performance of SM using numerical substrates of realistic WM generated with CATERPillar. By applying the SM to signals derived from these substrates, we gained insights into the mechanisms underlying changes in its estimates, as well as the conditions under which its estimations deviated from the GT.

### 4.1 Complex morphologies

In the SM, axons are idealised sticks with negligible width. The validity of this assumption had been verified in the human brain using very high diffusion weighting, which indeed yielded a dependence of the powder-average signal as 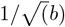 [41], in agreement with Callaghan’s model of randomly oriented sticks [82]. Nonetheless, it has not been established to what extent the effective GT diffusivities vary with morphological complexities such as undulations and beading, nor how biased SM estimates would be relative to the effective GT. Indeed, our results confirmed the sensitivity of SM estimates to undulation and beading.

Increasing axonal undulations resulted in a marked decrease in effective *p*_2_, *D*_*a*_, and *D*_*e*∥_. The orientation dispersion reflected by *p*_2_ was derived from water diffusion over approximately 50 ms, corresponding to a diffusion length of about 15 *µ*m. Because undulations can occur over such short length scales, both micro- and macroscopic orientation dispersion determine the total orientation dispersion [83], and higher undulation amplitudes were therefore expected to reduce *p*_2_. Similarly, *D*_*a*_ and *D*_*e*_∥ decreased because undulations hindered diffusion along the axonal axis. These findings are in agreement with a previous study [84] which shows the increase the estimated ODF and decrease *D*_*a*_ from the GT values with increasing undulation strength. In contrast, the increase observed in *D*_*e*⊥_ may arise from the formation of larger inter-axonal gaps oriented perpendicular to the fibre direction. Overall, although undulations influenced the effective GT values, the accuracy of the SM estimates with respect to them remained highly stable, with the exception of *D*_*a*_ which was mildly overestimated at high undulation coefficients.

Increasing the beading strength led to a decrease in the GT *p*_2_, *D*_*e*∥_ and especially *D*_*a*_ (which was similar to the effect observed with increasing undulation coefficients, but to a greater extent) and to an increase in *f* . The trends of the SM estimates followed those of their corresponding GTs, however, *f* and *D*_*a*_ were increasingly overestimated, whereas *p*_2_ was progressively underestimated at higher beading amplitudes. In healthy WM, the mean CV of axonal radius is typically around 0.2-0.3, as reported by [67, 68, 85]. At this level, the inaccuracies of these parameters were rather mild, indicating that *f, p*_2_, and *D*_*a*_ may not be too biased by beading under normal conditions. In contrast, as increased axonal beading has been reported in pathological tissues, the resulting overestimation of *f* and *D*_*a*_, and underestimation of *p*_2_, may be more problematic. Heavily beaded axons may indeed no longer be modelled as infinitely long sticks. The beading in pathology could go beyond the range investigated in this study, and thus introduce significant biases when interpreting microstructural alterations using the SM in these contexts.

The sensitivity of dMRI signals to axonal beading and undulations has been examined in several previous studies employing Monte Carlo simulations in numerical substrates [67, 68, 86–88], often with the goal of investigating their impact on axonal diameter estimation. These studies consistently reported decreases in mean diffusivity (MD) and apparent diffusion coefficient (ADC) with increasing undulation amplitude and beading strength. Further experimental investigations have confirmed that such morphological alterations occur due to axonal damage in pathological conditions such as spinal cord injury [89], multiple sclerosis [90–92], traumatic brain injury [93] and ischemic stroke [86], where reductions in parallel diffusivity have been proposed as a biomarker of axonal damage [68, 86, 89, 92, 94–96]. Our findings are therefore in strong agreement with these previous results, as both *D*_*a*_ and *D*_*e*_∥ (particularly *D*_*a*_) exhibited substantial decreases with increasing undulation and beading strength.

### 4.2 Axonal packing density

In this section, the substrates contained axons that were both beaded and undulating, with varying packing densities. Although different packing levels may have induced subtle differences in undulation amplitude and beading strength, we initially did not expect these variations to strongly affect estimation accuracy.

The accuracy and precision on *f* estimates were excellent throughout the range considered. Surprisingly though, our results revealed that for GT *f* values below 0.3, the model markedly underestimated *p*_2_. A smaller underestimation was also observed at GT *f* values above 0.5. According to [97], a more isotropic diffusion signal can arise either from an increase in fibre dispersion (lower *p*_2_) or from reduced anisotropy of the single-fibre response function (for example, due to low packing density or weaker diffusion restriction). It appears that, at low packing density, the model explained the increased isotropy primarily through greater fibre dispersion, leading to underestimated *p*_2_. This is likely due to the model design, where coherent fascicles made up of IAS and proximal EAS were convolved with the ODF. At low *f*, the EAS becomes much looser (*D*_*e*∥_ and *D*_*e*⊥_ nearly coincide) thus the medium may be interpreted as highly dispersed.

Overall, all estimates except *f* showed a bell-shaped dependence on fibre packing. For *D*_*a*_, the lowest and highest packing densities yielded the most accurate estimates, while intermediate packing led to overestimation. In contrast, the other parameters were most accurately estimated at intermediate packing densities.

In terms of effective GT, *D*_*a*_ was quite stable across substrates, slightly decreasing with higher axonal density, possibly due to altered beading and undulation to fit in more axons. The SM *D*_*a*_ estimate though showed a different pattern. In the noiseless case, it followed the bell shape previously described. In the finite SNR cases, the shape was monotonous, with best accuracy for high *f* and increasing overestimation with lower *f* . In the noiseless case, the bell shape may reflect some intrinsic spurious correlation between *p*_2_ and *D*_*a*_, although the latter was not identified in recent noise propagation work [98]. *D*_*a*_ is notoriously one of the most challenging parameter to estimate [11, 98] and its accuracy may further drop with low axon density, as its signature in the overall signal decreases.

### 4.3 Orientation dispersion

When assessing the impact of orientation dispersion on parameter estimation, low GT *p*_2_ values were associated with some overestimation of *f* but estimates remained otherwise accurate.

Regarding *D*_*a*_, a clearer picture emerges when considering fibre configuration. In voxels containing a single fibre population, *D*_*a*_ was overestimated in substrate E, where axons were highly aligned, but accurately estimated in substrate D, which featured axonal fanning modelled with a Watson distribution. This behaviour aligns with previous observations showing a negative relationship between axial diffusivity and ODF (i.e., a positive relationship between *D*_*a*_ and *p*_2_) for ODF values below 0.1, corresponding to well-aligned axons [84]. However, other work has reported the opposite trend (a positive relationship between axial diffusivity and ODF) when comparing SM estimates to electron-microscopy data [63]. Interestingly, this latter behaviour also emerged in our study when multiple fibre populations were introduced (substrates A–C), where *D*_*a*_ became increasingly overestimated as dispersion increased.

Because the SM assumes a single fibre population per voxel, voxels with high ODF values (indicating multiple fibre populations) are expected to yield larger estimation errors [99] and should therefore be interpreted with caution [84]. Using the ball-and-stick model, [100] investigated the effect of the inter-fibre angle on estimation accuracy and reported the most reliable reconstructions for crossing angles of *ϕ* = 45^°^–60^°^, with larger errors observed in voxels containing either highly aligned or widely crossing fibres. This observation aligns well with our findings, as the most accurate SM estimates were obtained in substrate D, which contained fanning axons.

Finally, when specifically examining voxels with two fibre populations (substrates B and C), *p*_2_ was overestimated. It is somewhat surprising that this effect was not also observed for three fibre populations in substrate A. One possible explanation is that, despite using isotropically distributed encoding directions, the orientations of the two fibre populations were not sufficiently sampled to ensure accurate estimation. Increasing both the number of diffusion-encoding directions [100] and the number of b-shells (beyond the five used in this study) [101] may improve the accuracy of SM parameter estimation.

### 4.4 Axonal permeability

When varying axonal permeability, the progressive underestimation of *f* with increasing permeability aligned with previous studies highlighting the impact of exchange on SM estimates in GM [31] and tumours [35]. The increase of *D*_*e*∥_ and *D*_*a*_ with permeability could be interpreted as reduced hindrance/restriction parallel to tortuous and beaded axons when membranes are permeable. Conversely, *D*_*e*⊥_ decreased with increasing permeability, as the radial diffusion of water molecules transitioning from the EAS to the IAS became more restricted. Though relatively minor, *p*_2_ gradually increased with axonal permeability.

Given the huge impact of permeability on the accuracy of *f* estimations, it is crucial to acknowledge that these estimates may not always be reliable if the impermeability assumption is not met. One might argue that the presented simulations lack realism, as myelin should be present to prevent exchange between compartments for all axons. However, histological studies [81] suggest that WM voxels typically contain a mix of myelinated and non-myelinated axons. Although myelinated axons comprise the majority, non-myelinated axons represent a considerable proportion and their prevalence increases in pathological conditions. Additionally, some WM voxels may contain other permeable structures such as dendrites and/or somas, for instance when GM and WM substrates mix. In order to effectively predict microstructural parameters in GM, models such as NEXI [31, 42, 102, 103] use the dMRI signal at multiple diffusion times to estimate the exchange time *t*_*ex*_ between the different compartments, linked to the permeability. However, NEXI is limited to GM and there is no study about the effect of different populations with different permeabilities on this parameter. An attempt has been made with eSANDIX [102] to describe a mixture of permeable and impermeable sticks but the model is very degenerate which compromises the precision of estimated parameters. Describing such substrates remains challenging as the minimal number of parameter required might be too high to avoid degeneracies with the available data.

It is important to note that the effect of increasing permeability on SM estimates observed in this analysis corresponded to a change in membrane permeability without geometrical or spatial occupancy changes. In other words, the effect does not reflect the scenario of myelin loss, which also translates into an increase in EAS. In studies investigating pathological conditions associated with myelin loss [4, 32, 104, 105], it was observed that *D*_*e*⊥_ (estimated using the SM, DKI or WMTI model) increased, contrasting with our simulations in this part. This suggests that the expansion of the EAS following myelin loss, which leads to an increase in *D*_*e*⊥_, has a more dominant effect than the increased permeability of demyelinated axons, which was found to decrease *D*_*e*⊥_ in our analysis.

### 4.5 Astrocyte permeability

Although the SM does not explicitly account for cellular components other than axons, it was expected to be sensitive to the presence of astrocytic processes and to potentially misclassify them as axons. This effect was indeed observed, particularly when astrocytes were impermeable. At zero permeability, *f* was overestimated compared to its GT (which considered only axons), as anticipated, with estimated values even slightly exceeding AVF+ASVF, which incorporated both axonal and astrocytic volume fractions. Furthermore, *p*_2_ was underestimated relative to its GT (which again considered only axons), reflecting the greater orientation dispersion of astrocytic processes compared with axons. At low GT *p*_2_ values, our previous investigation has shown that *f* tends to be overestimated, which may further explain why the estimated *f* exceeded the AVF+ASVF in this case. As the permeability of astrocytes increased, astrocytic water gradually became classified as extra-axonal space and the estimated axonal *f* and *p*_2_ converged towards their GT values.

Compartment-specific diffusivity estimates were relatively stable over astrocyte permeability values. In the noiseless case, one noticeable trend was a gradual increase in *D*_*a*_ and decrease in *D*_*e*∥_ and *D*_*e*⊥_, which can be explained by the shift of astrocytic water from being associated with the axonal compartment (permeability=0) to being associated with the extracellular water (permeability = 10^−4^ m/s). Indeed, as astrocytic processes are shorter, branched and overall less straight than axons, their association with axonal water would decrease *D*_*a*_. Conversely, when they contribute to the extra-axonal compartment, they make for a more tortuous medium, hence a decrease in the apparent *D*_*e*∥_ and *D*_*e*⊥_. This influence is attenuated by the addition of noise. In contrast, the GT *D*_∥_ and *D*_⊥_ were overall stable, confirming that the changes in compartment diffusivities are driven by shifts between which compartment the astrocytes are assimilated to, not affecting much the weighted average of axial or radial diffusivity in the voxel.

These observations are partially consistent with previous findings by [106], who reported a marked reduction in ADC following the knockdown of aquaporin-4 water channels, which decreased astrocytic permeability. As astrocyte morphology was assumed to remain unchanged in that study, the reduction in ADC was attributed solely to the decrease in permeability, indicating that higher permeability facilitates overall water mobility. In our simulations, the slight increase in *D*_∥_ and *D*_⊥_ with lower astrocyte permeability was considerably smaller than what was observed by [106]. This attenuated response may result from the relatively low astrocytic volume fraction in our substrates, though they were reported to be as such in deep WM (1% soma volume fraction and 6% processes volume fraction [81]). In [106], the data came from rodent GM, in which the density of astrocytes may have been higher.

Overall, as astrocytes in physiological conditions are known to be quite permeable, these results confirm typical SM hypotheses, where the “extra-axonal space” is assumed to lump contributions from actual extra-cellular space and other non-axonal structures, such as astrocytes, microglia, etc [20]. As a result, this suggests that the SM extra-axonal compartment may absorb contributions from glial cells, and EAS diffusivities could therefore be sensitive to glial-related alterations. This is in line with previous works that linked EAS diffusivities to neuroinflammation and to changes in glial density in WM [32, 92, 107–109].

### 4.6 Myelin Volume fraction

The inclusion of myelin was also expected to challenge the SM estimations, as this compartment is not explicitly represented in the model. Since myelin is assumed to not contribute to the overall MRI signal due to the short *T*_2_ of myelin water, it is essential to distinguish between *f*, which accounts only for signal-bearing compartments, and the AVF+MVF, which reflects the total intracellular volume, as previously discussed. When the MVF is properly factored out in defining the effective GT, all SM parameter estimates remained relatively close to their GT values in our results. An underestimation of *p*_2_ and overestimation of *D*_*a*_ were observed, likely driven by the low packing density (see corresponding discussion above) rather than the presence of myelin itself. The estimated *f* lay between the AVF (computed without accounting for myelin) and the AVF+MVF, as expected, although it was slightly overestimated compared to its GT.

Regarding the diffusivity estimates, a decrease in both *D*_*e*∥_ and especially *D*_*e*⊥_ (both the effective GTs and their estimates) could be observed, whereas *D*_*a*_ was rather constant. The large decrease in *D*_*e*⊥_ with higher MVF was expected, as this variable has been considered the main biomarker for myelination and demyelination in countless studies [4, 32, 92, 95, 105, 110–115]. Myelination occurs in early brain development and later plasticity, while demyelination occurs in aging or pathologies such as multiple sclerosis, Alzheimer’s disease, schizophrenia and traumatic brain injury.

### 4.7 Diffusion Time

Increasing the diffusion time improved the accuracy of morphological parameter estimates (*f* and *p*_2_) in our simulations, thanks to the coarse-graining effect. A convergence of the diffusivity estimates towards their GT *D*_∞_ was expected due to the time dependence of diffusivity in substrates complex enough to have short-range disorder characteristics [6, 34, 116], particularly in the axial diffusivities, where the typical length scale of axonal irregularities matches the diffusion length probed here, as also reported experimentally [104]. It is important to note, however, that *T*_2_ relaxation was not accounted for in our simulations and not predicted by the SM (though an extension of the SM could do so [5, 101, 117]). In practice, the SNR decreases with longer diffusion times, which can in turn reduce estimation accuracy. Therefore, an optimal balance must be achieved, where water molecules probe the substrate for a sufficiently long time while maintaining an adequate SNR. Previous work optimised SM protocols with diffusion times between 92 and 122 ms [5].

### 4.8 Noise

Across all figures comparing results with and without noise, we observed that decreasing SNR led not only to an increased standard deviation (reflecting lower precision) for all parameter estimates, but also to consistent biases, most notably in *p*_2_ and *D*_*e*∥_. At lower SNRs, *p*_2_ was systematically underestimated, while *D*_*e*∥_ was consistently overestimated. In contrast, the parameter *f* remained largely unbiased across all SNR levels. As reported by [14], this behaviour is expected: volume fractions can be estimated reliably from high-quality data, whereas diffusivity estimates tend to be noisier and more biased across acquisition schemes. Furthermore, [5] identified *D*_*e*∥_ as the most noise-sensitive parameter, being the most difficult to estimate and therefore the most influenced by training data. Despite these biases, the same study concluded that the SM remains sufficiently robust to capture biological variability across WM regions (and potentially pathological variability) making it “good enough” for practical use. Regarding *p*_2_, our findings are consistent with prior work showing that low SNR values (below 15) substantially hinder the accurate estimation of the fanning extent [24]. Although our simulations only considered SNR levels of 50 and 100, a clear bias in *p*_2_ was already evident, suggesting that this effect would likely be even more pronounced at lower SNRs (typically close to 30 in WM).

Our analysis thus extends previous work by explicitly revealing the direction of the bias: *p*_2_ tends to be underestimated, while *D*_*e*∥_ is overestimated in the presence of noise. The noise introduced in this study followed a Rician distribution, as the magnitude data contained independent noise components in both the real and imaginary channels.

### 4.9 Summary

Overall, the relative changes in SM estimates when varying parameters (summarised in Figure 12A) indicate that:

1. The intra-axonal volume fraction *f* (both effective GT and its SM estimation) increased as expected with stronger beading, higher axonal packing density, and greater myelin volume. Its estimation was overestimated with stronger beading and high axonal dispersion, underestimated with high axonal permeability.
2. The effective orientation dispersion index *p*_2_ was influenced by undulation and beading. Its estimation via the SM was quite biased by axonal packing density in particular, but also by beading and when astrocytes were impermeable.
3. The effective intra-axonal diffusivity *D*_*a*_ was closely tied to axonal morphology, decreasing with stronger beading and undulation. Furthermore, it became overestimated in the SM in most conditions.
4. The effective parallel extra-axonal diffusivity *D*_*e*∥_ was strongly affected by axonal permeability, but also by packing density, dispersion, and undulation strength. Its estimate was also the most noise-sensitive parameter, showing systematic overestimation at low SNR levels (SNR = 50). At higher SNR levels, it was consistently accurately estimated.
5. The effective perpendicular extra-axonal diffusivity *D*_*e*⊥_ was mainly sensitive to axonal packing density, axonal permeability, and myelin volume fraction, making it a potential biomarker for (de)myelination, but also to undulation. Its SM estimate was also consistently accurate with respect to the effective GT.

The most unexpected findings of this study were:

1. the bell-shaped behaviour of all estimates except *f* with increasing axonal volume fraction *f*,
2. the overestimation of *p*_2_ occurring only in the presence of two crossing fibre populations,
3. the strong overestimation of *D*_*a*_ when multiple crossing populations were present. Future work will focus on understanding the origin of these behaviours in the SM estimates.

## 5 Limitations

Monte Carlo simulations were conducted within voxels measuring 150 *µm* in width, with walkers initialised inside a central cube of 110 *µm*. While this configuration reduces the likelihood of walkers reaching the voxel boundaries, it does not fully eliminate it. Encounters with the mirror boundaries can induce artificial turning behaviour, potentially affecting the *p*_2_ estimation.

Additionally, though the impact of each variable on the SM estimates was assessed, we did not investigate how they impact them jointly. For example, cell permeability may affect the results even more when the packing density is higher. Future work could attempt to assess the combined effects of these parameters to better capture their interdependencies.

## 6 Conclusions and Future Works

Using numerical WM substrates, we were able to assess the variation of the SM parameter estimations across various substrates, and to identify the conditions to which the model parameters were most sensitive, but also the ones under which the parameter estimates exhibited the greatest biases. Awareness of potential biases under different conditions is crucial for a reliable interpretation of results in applications to real tissue. The findings revealed that among others, axon permeability, high orientation dispersion and low axonal packing density considerably affected the accuracy of the SM estimations. All of these conditions considerably affected the estimation of *f, p*_2_ and *D*_*a*_. SM-derived EAS compartment diffusivities on the other hand displayed an excellent accuracy in the conditions of our simulations. However, *D*_*e*∥_ was particularly biased with the addition of Rician noise.

In future work, we plan to further enhance the realism of our numerical substrates by incorporating blood vessels in CATERPillar. An updated version of the MC/DC tool with integrated *T*_2_ relaxation could then be used to assess how vasculature-induced inhomogeneities influence diffusion-weighted signals.

## 8 Code availability statement

The source codes for both the CATERPillar tool and the modified MC/DC tool have been made publicly available on GitHub: https://github.com/jazz031195/CATERPillar and https://github.com/jazz031195/Permeable_MCDS.

## 9 Author Contribution

Conceptualization: I.J, J.N.D; Methodology : I.J, J.N.D; Validation : J.N.D; Formal analysis: J.N.D; Writing-original draft: J.N.D; Writing - Review & Editing: I.J, J.N.D, Q.U, J.R.P; Visualization: J.N.D; Supervision: I.J.;

## 10 Acknowledgements

This work was supported by the Swiss Secretariat for Research and Innovation (SERI) under an ERC Starting Grant award ‘FIREPATH’ MB22.00032, and by Swiss National Science Foundation (SNSF) grants no. 194260 and 10000465.

