## Supplementary material for "Validating the Standard Model of diffusion MRI in white matter with Numerical Substrates": S1

### Supplementary Materials

#### 1 Assessing the impact of the number of random walkers in MC/DC

To verify that a density of one random walker per $\mu m^3$ (corresponding to a total of 1,330,000 walkers) yields stable and accurate MRI signal estimates, we evaluated the coefficient of variation ($\mathrm{CV} = \frac{\mathrm{std}}{\mathrm{mean}}$) of the signal across several walker densities. Because a lower CV indicates greater signal stability, our goal was to determine the density at which the CV plateaus, thereby justifying our choice. To this end, we performed the Monte Carlo simulations on a substrate containing axons with a packing density of 0.5.

We performed 40 independent Monte Carlo simulations, each with 133,000 walkers within the IAS and EAS compartments separately. The signal from these independent runs were normalised and then combined in random groups to emulate higher walker densities.

We compared the signal obtained when combining 1, 2, 5, 7, 10, 15, 20, and 30 runs with 133,000 walkers each. For each group, the signal was generated 10 times with random sampling (with replacement), and the mean and CV across these 10 groups.

This procedure was repeated for all walker densities, and the resulting mean and CV values were averaged over 100 repetitions. The final results (Figure S1) present the mean powder-averaged DWI signal and corresponding CV as functions of walker density and $b$-value. At a density of 1 walker/$\mu m^3$, the CV of the normalised powder-averaged signal was below 0.002 for the IAS and 0.02 for the EAS. The highest CV in the EAS occurred at a $b$-value of 5 ms/$\mu m^2$, consistent with expectations since such high $b$-values are known to attenuate the extracellular signal. The chosen density is marked by an arrow in Figure S1. The elbow point (identified using the KneeLocator library [1]) is located at 0.70 walkers/$\mu m^3$, just before the arrow. In optimisation problems, the elbow typically marks the point beyond which further increases are not worth the additional cost. Selecting the next density step therefore provides a small safety margin. Thus, our results indicate that choosing one walker per $\mu m^3$ was appropriate, as the elbow occurs immediately before this value.

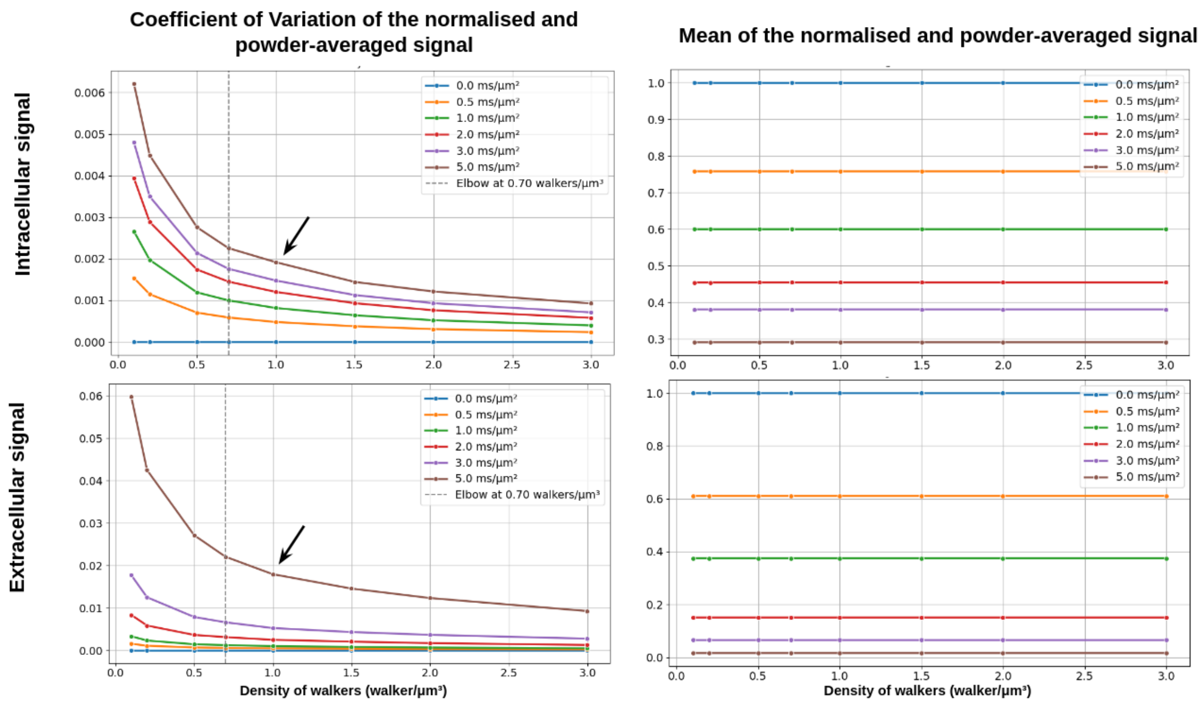


Figure S1: Average coefficient of variation (CV) and mean normalized powder-averaged signal. Mean values and CVs were computed in groups of 10 and recalculated 100 times through resampling. Results are shown for multiple $b$-values (indicated in the legend) in ms/$\mu$m$^2$. The arrow marks the walker density selected for the Monte Carlo simulations in this study.
